# A novel pH-sensitive probe to quantify autophagy on high throughput/content imaging platforms

**DOI:** 10.1101/2025.05.13.653902

**Authors:** GD Ciccotosto, M Jana, L Galluzzi, L Mitchell, P Gupta, SGB Furness, M Lazarou, DL Hare, PJ Wookey

**Author notes:** Corresponding author Assoc/Prof Peter J Wookey, Department of Medicine-Austin, University of Melbourne, Austin health, Level 7, Lance Townsend Building, Studley Road, Heidelberg, Victoria 3084, Australia.

## Abstract

Autophagy is a highly-conserved mechanism that ensures the lysosomal degradation of disposable or potential toxic cytosolic entities in support of cell survival and adaptation to stress. Autophagy is deregulated in various pathological conditions including cardiovascular, neurological, neoplastic-autoimmune and degenerative disorders. However, clinically relevant pharmacological modulators of autophagy remain elusive, calling for the development of novel screening approaches that are amenable to high throughput/content applications. Here, we describe a simple method to detect autophagy in cultured mammalian cells based on the pH-sensitive fluorescent conjugate CalRexin™:pHrodo™ Red and high throughput/content imaging. CalRexin™:pHrodo™ Red is promptly taken up by endosome-amphisome-autolysosome (low pH ∼4.5) pathway, which is connected to canonical autophagy, culminating with a bright red fluorescence. Importantly, this system allows for the discrimination between autophagic responses coupled with the endosome-amphisome-autolysosome pathways and the endosome-lysosome pathway as elicited by receptor-driven endocytosis and phagocytosis. In human cervical carcinoma sHeLa cells, the accumulation of CalRexin™:pHrodo™ Red as elicited by canonical autophagy activators like the mTOR inhibitor rapamycin or serum starvation was suppressed by conventional inhibitors 3-methyladenine or chloroquine. Similarly, *RB1CC1-/-(FIP200)* as well as *ATG5-/-* sHeLa cells (which bear genetic defects in two different steps of autophagy) were unable to accumulate CalRexin™:pHrodo™ Red upon exposure to autophagy activators. Thus, CalRexin™:pHrodo™ Red provides a novel approach for imaging autophagy on high throughout/content platforms to screen large libraries for the identification of novel autophagy-targeting agents.

## Introduction

Macro-autophagy (herein autophagy) is a highly conserved mechanism through which all eukaryotes dispose of dysfunctional, supernumerary or potentially toxic cytosolic entities upon inclusion in a double-membraned organelle commonly known as autophagosome followed by lysosomal degradation [1,2]. Besides operating at baseline levels in healthy cells, such a process can be activated by a variety of stressors including, but not limited to, cytotoxic agents, infective challenges and metabolic alterations [3]. Abundant preclinical and clinical literature demonstrates that autophagy is critical not only for embryonic and post-embryonic development, but also for the preservation of adult tissue homeostasis [4]. Moreover, autophagy defects appear to aggravate disease progression in a variety of pathological conditions, including (but not limited to) cardiovascular, neurological, neoplastic-autoimmune and degenerative disorders [5-9].

Autophagosomes are formed by the elongation and maturation of single-membraned “phagophores”, which appear to emerge from specialized domains of the endoplasmic reticulum, a process that is finely regulated by more than 100 proteins [10]. Of note, most of these proteins have autophagy-unrelated functions, often (but not always) linked to vesicular trafficking [11]. Autophagosome elongation culminates with their closure around autophagy substrates (e.g., permeabilized mitochondria), resulting in mature double-membraned autophagosomes that can fuse with late endosomes (pH ∼ 5.5) [12] by a RAB11A, member RAS oncogene family (RAB11A)-dependent pathway [13,14], to form transient amphisomes [12]. Ultimately, the latter fuse with the lysosomes (pH ∼4.5) to generate autolysosomes, culminating with the pH-dependent activation of hydrolases and degradation of autophagy substrates [10].

Despite such a broad role in disease [10], currently available pharmacological modulators of autophagy of clinical relevance are scarce and largely unspecific [15,16], encompassing the mechanistic target of rapamycin complex 1 (mTORC1) inhibitor rapamycin (which activates autophagy but also has major effects on proliferation) as well as the lysosomal inhibitors chloroquine and hydroxychloroquine (which inhibit autophagy but also other lysosome-dependent pathways) [15,16]. Thus, considerable screening efforts are being devoted to the discovery of novel autophagy modulators. However, current methods to investigate autophagic responses are either incompatible with high throughput/content platforms (e.g., the immunoblotting-based assessment of autophagosome maturation markers), or they quantify the size of specific autophagy compartments but not true autophagic flux (e.g., the fluorescence-based assessment of autophagosome formation) [2,17,18].

We devised a novel strategy that quantifies *bona fide* autophagic responses based on the pH-sensitive probe CalRexin™:pHrodo™ Red, an antibody conjugate that selectively binds to calcitonin receptor (CT Receptor) and – in the context of increased autophagic flux – is internalized via early endosomes (pH ∼6) [19,20] that [1] mature into late endosomes (pH ∼5.5) [21,22], [2] fuse with autophagosomes to form the transient amphisomes, and [3] reach the lysosomal compartment (pH ∼ 4.5). The latter enables CalRexin™:pHrodo™ Red to acquire a bright red fluorescence. We chose this receptor because CT Receptor appears to be associated with inflammatory signaling [23], and pre-apoptotic cell stress [24], and the maintenance of quiescence in adult muscle satellite stems cells [25-28], three settings that are associated with autophagy activation [4,29,30].

Here, we provide experimental validation to CalRexin™:pHrodo™ Red as a suitable probe to quantify autophagic responses in high throughput/content imaging platforms based on cultured cells exposed to activators of autophagy alone or in the context of pharmacological or genetic autophagy inhibition.

## Materials and Methods

### Cell lines

The human cervical carcinoma Hela (sHela) cell lines, including wild-type [WT], the knockout [KO] *Atg5* (lacks lipidation of LC3B) [31-33] and *RB1CC1*^*-*^(*FIP200*) sHeLa cell lines [34], were cultured in DMEM containing high glucose and sodium pyruvate (Thermo Fisher Scientific [TFS, 11995-065]), and supplemented with 10% foetal bovine serum (FBS, [TFS, 10100147]), 2mM glutamax (TFS, 35050061), MEM non-essential amino acids (TFS, 1114005), 100mM HEPES (Sigma, H3375) pH 7.4, 10 U/mL penicillin-streptomycin (TFS, 15140122).

Cell lines were maintained in a humidified incubator (5% CO_2_, 37 ^o^C) with media changes or cell passaging performed at a confluency of 80%. Cells were dislodged with Trypsin/EDTA (TFS, 15400054), harvested, pelleted and a single cell suspension prepared and the number of cells counted. For Operetta experiments cells were seeded at a density of 10,000 cells per well in a sterile 96 well, black-walled, optically clear cyclic olefin flat-bottom plate, (Cell Carrier-96 Ultra [PerkinElmer, 6055302]) using only the inner 60 wells of the plate while the outer wells were supplemented with 1x sterile PBS buffer to minimize evaporation from the experimental wells. Plate surfaces were precoated with Matrigel solution ([Corning, 354230] diluted 1:100 in cold PBS, 100 μL per well) overnight at 37 °C.

For confocal microscopy cells were seeded at a density of 20,000 cells per well in μ-Slide 8 well (DKSH, 80821) slides precoated with Matrigel solution. Cells were allowed to adhere to surface overnight before treatment and imaging. Experiments were performed using cell lines passaged less than 20 times

Depleted FBS (dFBS) was prepared by ultracentrifugation of FBS (TFS, 10100147) at 100,000 g for 24 hr at 4 °C and the clear supernatant devoid of exosomes, (top 30% of supernatant) was collected as dFBS. Exosome free FBS (eFBS) was sourced commercially (TFS, A2720801).

Rapamycin (Sigma, R0395) and serum starvation were used to induce autophagy while 50 μM chloroquine (Sigma, C6628) or 10 mM 3-methyladenine (Sigma, M9281) inhibit autophagy.

### High Content Screening

Cells were seeded as described above the night before and were imaged on an Operetta high-content screening system (Operetta CLS) operated by the Harmony software (Harmony version 4.1). At T=0 h, the nuclear dye Hoechst 33342 (Sigma), CalRexin™:pHrodo™ Red (4 μg/mL, Apop Biosciences Pty Ltd, www.apopbiosciences.com), CYTO-ID® (ENZO, ENZ-KIT 175-0500), LysoTracker™ Green (300ng/mL TFS #L7526) were mixed with fresh media and added to the cells. Cells were imaged hourly over a 24 h period with images set to non-confocal, being acquired at a single focal plane every hour using a 40x high Numerical Aperture objective. The system was maintained at a constant temperature of 37 °C and 5 % CO_2_ for optimal cell growth. The Operetta platform is programmed to collect images (40x magnification) from 60 central wells of a 96-well optical plate. From each imaged well, at least five field positions were chosen (four from the periphery and one central) which results in analysis of approximately 100 cells per well. Each experimental well is repeated in triplicate in each plate.

sHeLa cells were transfected using several different conditions to maximise the transfection frequency with the vectors expressing RFP-GFP-LC3BII according to the manufacturer’s instructions and as supplied in the Premo™ tandem sensor kit (TFS, P36239). The optimal rapamycin concentration for induction of autophagy is cell line dependent and for sHela cells 8 μM was used (see dose-response data in Figure 1). The inhibitors chloroquine (50 μM) and 3-methyladenine (10mM) were used at recommended concentrations.

**Figure 1.**
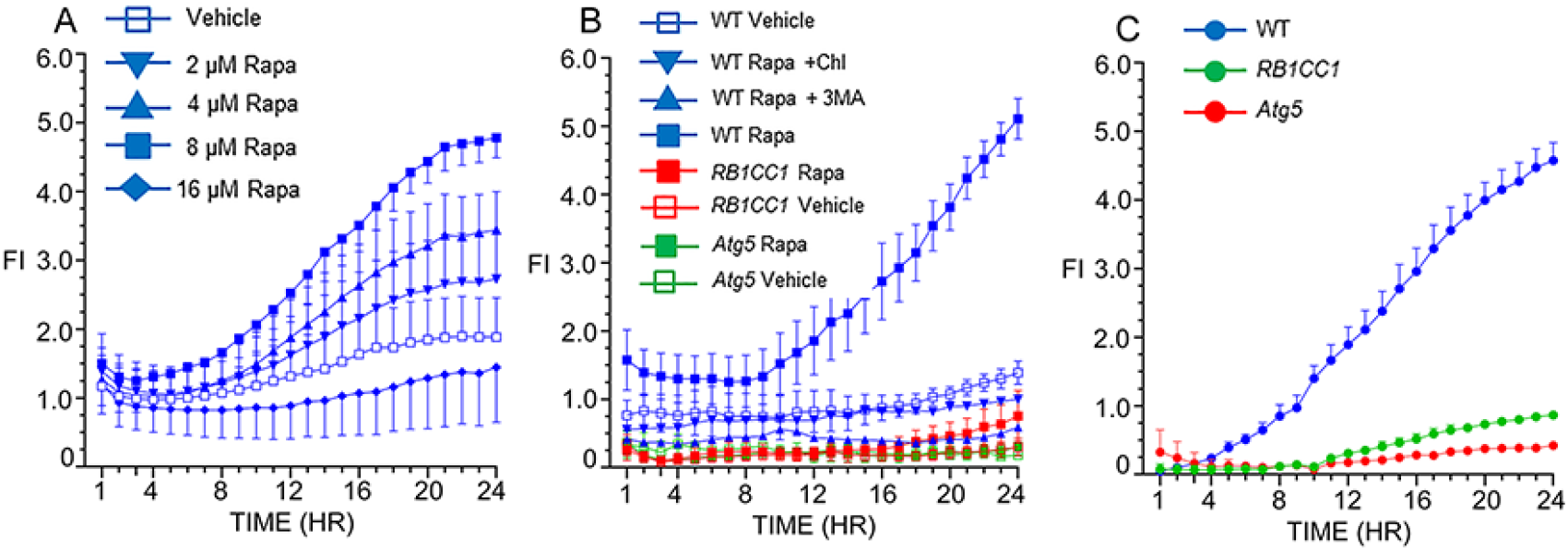
Operetta profile of fluorescence intensity of intracellular CalRexin™:pHrodo Red (4 μg/mL) in live sHeLa wildtype [WT], sHeLa *RB1CC1* and sHeLa *Atg5* cells with induced autophagy with 8 μM rapamycin versus vehicle controls imaged hourly for 24 hours. Each data point is the mean of three independent experiments +/-SEM. **Figure 1A**. Increased fluorescence of sHeLa cells in response to the dose response of rapamycin: vehicle, blue open squares; 2 μM, blue inverted triangles; 4 μM, blue triangles; 8 μM, closed squares; 16 μM, closed diamonds. **Figure 1B**. Increased fluorescence of sHeLa cells in response to: 8 μM rapamycin, blue closed squares; vehicle, blue open squares; 50 μM chloroquine with 8 μM rapamycin, closed blue inverted triangles; 10 mM 3-methyladenine with 8 μM rapamycin, closed blue triangles; sHela *RB1CC1* with 8 μM rapamycin, closed red squares,; sHela *RB1CC1*, vehicle, open red squares; sHela *Atg5* with 8 μM rapamycin, closed green squares; sHela *Atg5*, vehicle, open green squares,. **Figure 1C**. Cells grown in the absence of serum (serum-starved without rapamycin): sHeLa WT, closed blue circles; sHeLa *RB1CC1*, closed green circles; sHeLa *Atg5*, closed red circles.

The following filter sets were used for imaging each dye: for Hoechst 33342, excitation was 360-400 nm and emission was 410-480 nm, 80 ms; for CalRexin™:pHrodo™ Red and RFP-LC3B, excitation was 520-550 nm and emission was 560-630 nm, 800 ms; for Lysotracker™ Green, CYTO-ID®, and GFP-LC3B, excitation was at 460-490 nm and emission was 500-550 nm, 300 ms. Excitation was provided by a Xenon lamp and transmission was set to 50 %. The brightfield channel was acquired with an exposure of 50 ms.

Cell autofluorescence was evident especially during the first few hours in the red channel used for imaging the CalRexin™:pHrodo™ Red and varied (1-3 FI absorbance units) between experiments. It was evident in the absence of CalRexin™:pHrodo™ Red and was subtracted from test time-points in which there was significant autofluorescence. For each experimental plate, each data point was determined in triplicate wells and within each well at least four field locations were selected for imaging (middle position and at least three peripheral sites) and these parameters were identical for each timepoint examined. These parameters ensured the presence of approximately 100 cells per well for data analysis.

For each experiment the data points were normalized between experimental repeats by subtracting autofluorescence values from raw value.

### Fluorescence Intensity quantitation

Time-lapse videos were created and annotated using FIJI image analysis software, version 1.53c (Fiji, RRID:SCR_002285). The average fluorescence intensity (FI) for these cells per well at each timepoint was measured using FIJI software to quantify average FI. Data from each well at time T represents one data point. To calculate the average FI, cell images were taken under identical exposure settings using the Harmony software settings and a panel of images including each time point was saved as Tiff files and imported into Fiji Image J software (NIH, v1.53r) for image analysis. The region of interest for each image was selected, covered the whole field, which were kept identical across all time points examined and the fluorescence intensity was measured for each field in an identical manner. Experimental samples were tested across triplicate wells and from each well, at least four field images were analysed. The calculated results represent the fluorescence intensity average of the fields in triplicate wells. Fluorescence intensity measurements are imported into a Microsoft Excel spreadsheet for further calculations and final graphs were prepared using Prism Graphpad (version 9.5.1). Experiments were repeated at least in triplicate and the error bars represent +/-SEM.

### Live Cell Imaging for Confocal Microscopy

Cells were grown and treated in phenol red-free culture media with 5% FBS in 8 well chamber slides and placed in an environmental chamber (Oko Labs) to maintain cells at a constant 37 °C with humidified 5% CO_2_ for optimal cell growth during imaging. Images were acquired using a Nikon CSA-W1 SoRa spinning disk confocal microscope (Nikon, Yokogawa Electric Corporation, Tokyo, Japan) at the Peter MacCallum Core Facility (Parkville, Australia), using the Nikon Ti-E, Yokogawa W1 spinning disk (50 μm) with a plan Apo λ 60x (NA 1.4) (RI 1.51). Images were collected using z-stacks of multi-channel fluorescence and image detectors and a 2x Andor Sona sCMOS camera, all operated by the NIS-Elements Software (NIS-Elements version 4.51). The following filter sets were used for imaging each dye: Hoechst 33342 was excited by a 405nm laser and emission filtered by 447nm, CalRexin™:pHrodo™ Red and RFP-LC3B (red channel) were excited by a 561 nm laser and emission filtered at 617 nm, Lysotracker™ Green, CYTO-ID® and GFP-LC3B dyes were excited by a 488 nm laser and emission filtered at 525nm. The brightfield images were acquired in the spinning disk mode with transmission set at 607nm and a 50 ms exposure. Image z-stacks were taken with 0.2 μm distance between slices and deconvoluted using the automatic mode in the NIS-Elements software. For negative controls, WT sHeLA cells were treated with 8 μM rapamycin for 24h and stained with Hoechst 33342 only to define the negative fluorescent signals for both the red and green channels.

### Conjugations of antibody CalRexin™

The mouse monoclonal antibody mAb2C4 (clone 46/08-2C4-2-2-4, IgG1 kappa, CalRexin™ [RRID: AB_2893093]) was developed using standard techniques described previously [35]. The immunization of mice and purification were performed in the dedicated laboratories at the Antibody Facility of the Walter & Elisa Hall Institute of Medical Research (Bundoora, Victoria, Australia) as the third-party provider. CalRexin™ binds within the N-terminal domain of the CT Receptor isoforms, CT_a_ or CT_b_ Receptor. The strategy to develop CalRexin™ has been described [35]. Data supporting the validation of the antibody CalRexin™ have been presented in the supplementary materials linked to Furness *et al*. [24]. CalRexin™ is a proprietary antibody of Apop Biosciences Pty Ltd (Melbourne, Australia: www.apopbiosciences.com).

For synthesis of the initial batch of CalRexin™:pHrodo™ Red (RRID: AB_2893121), the method for conjugation of CalRexin™ (mAb2C4) with pHrodo™ Red NHS esters ([TFS, P36600], MW 650) has been described previously [35]. The molar ratio of antibody to dye was 1:15 and 15 mg of antibody was conjugated with 1 mg of pHrodo™ Red NHS esters. The antibody was desalted by centrifugation using 5mL Zeba column (TFS) and the buffer replaced with 100mM sodium bicarbonate, pH 8.3. The dye pHrodo™ Red NHS esters (MW 650, 1 mg, 1.54 μmoles) was dissolved in 150 μL of anhydrous DMSO and added dropwise to the antibody in bicarbonate buffer. After 1 h gentle inversion at RT the reaction was quenched with 50 μL of 1.5 M triethanolamine (20% solution in bicarbonate buffer) for 3 h. The conjugate was separated from unreacted and other small molecules on a Zeba column and the buffer was exchanged for PBS. The final concentration was 1 mg/mL with a degree of labelling [DoL] of 1.7.

For subsequent batches of CalRexin™:pHrodo™ Red and prior to conjugation, CalRexin™ was buffer exchanged into 20 mM bicine, 150 mM NaCl pH 8.3 using a HiPrep 26/10 Desalting column (Cytiva). Reaction was performed with 8 equivalents of pHrodo™ Red NHS esters ([TFS, P36600], MW 650) using a 2.6 mM solution in anhydrous DMSO or with 8 equivalents of pHrodo™ Red iFL STP ester (DoL 1.9, MW ∼ 1000, [TFS, P36010]) using a 5.7 mM solution in anhydrous DMSO and incubated overnight at RT. The reaction was then quenched with a 150-fold molar excess of ethanolamine and incubated for 3 h at RT.

Immunoaffinity chromatography was performed using an Econo-column® (Bio-Rad) packed with CalRexin™ peptide ligand affinity resin. The acetamidomethyl (Acm) capped CalRexin™ peptide ligand (CSGSPSEKVTKYC(Acm)DEKGVWFK, Auspep Australia) was conjugated to Epoxy Sepharose 6B (Cytiva) resin according to the manufacturer’s protocol. The affinity resin was first equilibrated with 5 column volumes (CV) of PBS, then the CalRexin™:pHrodo™ Red reaction mixture was added dropwise. The resin was washed with 5 CV of PBS then eluted with elution buffer (0.1 M citrate, 0.15 M NaCl, 2 M arginine, pH 3.0) and neutralized with 3 M Tris pH 8.1.

Further purification of CalRexin™:pHrodo™ Red conjugates was carried out by gel filtration using a HiLoad 26/600 Superdex 200 pg (Cytiva) equilibrated in PBS. CalRexin™:pHrodo™ Red conjugates were filtered with Acrodisc® 0.2 μm syringe filter and the DoL determined spectroscopically to be 1.1 and 1.0 for pHrodo™ Red iFL STP and pHrodo™ Red NHS conjugates, respectively. To synthesise the isotype control, mouse IgG1 antibody (BioLegend, 400102, clone MOPC-21) was conjugated with pHrodo™ red iFL STP ester ([TFS, #P36010], MW ∼1000) and the resulting DoL was 1.35.

### Statistics

Graphical presentation and statistical analysis were performed using Prism 9.5.1 software (GraphPad Software, Inc., San Diego, CA). Data are represented as mean +/-SEM.

## Results

To elucidate the ability of CalRexin™:pHrodo™ Red to measure autophagic responses, we exposed human cervical carcinoma sHeLa, human osteosarcoma MG63 and immortalized mouse fibroblast NIH-3T3 cells to increasing concentrations of rapamycin, together with staining using 4 μg/mL CalRexin™:pHrodo™ Red and imaging hourly for 24 h. Importantly, CalRexin™:pHrodo™ Red exhibits a bright red fluorescence at pH of 5 or lower (Figure S1A). Rapamycin elicited a time- and dose-dependent increase in CalRexin™:pHrodo™ Red fluorescence in sHeLa cells (Figure 1A). Similar results (but with different kinetics and magnitudes) were obtained with MG63 cells (Figure S1B). As a control of specificity, 8 μM rapamycin was unable to elicit an increase in red fluorescence when CalRexin™:pHrodo™ Red was replaced by the isotype control IgG1:pHrodo™ Red (Figure S1C). Moreover, the performance of CalRexin™:pHrodo™ Red synthesized as the NHS ester or an iFL STP ester was fairly comparable, both with a degree of labeling of 1.9 and the NHS ester (MW ∼650 daltons, extinction coefficient 65,000) exhibiting slightly improved maximum fluorescence over 24 h as compared to the iFL STP ester (MW∼ 1000 daltons, extinction coefficient 65,000) (Figure S1D).

Importantly, the increase in CalRexin™:pHrodo™ Red fluorescence as elicited by 8 μM rapamycin was abrogated by the lysosomal inhibitor chloroquine as well as by 3-methyladenine (Figure 1B), an experimental inhibitor of autophagy that operates at early steps of the process. Moreover, the uptake of CalRexin™:pHrodo™ Red by sHeLa cells and consequent red fluorescence as promoted by 8 μM rapamycin, was abrogated by the deletion of RB1 inducible coiled-coil 1 (*RB1CC1, FIP200*) or autophagy related 5 (*ATG5*) (Figure 1B), which code for two key components of the autophagy machinery [10]. Similar results were obtained when serum deprivation was employed to elicit autophagy in sHeLa cells rather than rapamycin (Figure 1C).

Interestingly, sHeLa cells cultured with exosome-depleted (d) or exosome-free (e) FBS underwent mild autophagic responses as detected by imaging with CalRexin™:pHrodo™ Red (Figure S2A). Those were lower in magnitude than autophagy elicited by serum deprivation (Figure S2A), but equally sensitive to RB1CC1 and ATG5 deletion (Figure S2B & C).

To ascertain the performance of CalRexin™:pHrodo™ Red as compared to other, commercially available reagents for the detection of autophagy, we transfected wild-type sHela cells with the Premo™ Autophagy Tandem Sensor, which encode the autophagy adaptor microtubule associated protein 1 light chain 3 beta (MAP1LC3B, best known as LC3B) in tandem with acid-sensitive GFP as well as acid-insensitive RFP, enabling the visualization of autophagosomes in yellow and lysosomes in red [36]. At 1 h incubation with 8 μM rapamycin, sHeLa cells expressing RFP-GFP-LC3B exhibited a bright red fluorescence (Figure S3C), confirming high transfection rates but this rapidly declined (potentially due to photobleaching) [37] and virtually no green fluorescence (Figure S3A & B). Twenty-four hours later, a few cells remained that exhibited bright yellow puncta (Figure S3D & E), for reasons that are not clear. As successful imaging on the Operetta platform depends on a large proportion of fluorescent cells the RFP-GFP-LC3B reporter does not seem ideally suitable for assessing autophagy on this (or similar) imaging platform(s).

Along similar lines, we assessed the performance of CYTO-ID®, a green dye that mainly binds phagophores and autophagosomes [38]. sHeLa cells exposed to 8 μM rapamycin and stained with both CYTO-ID® and CalRexin™:pHrodo™ Red exhibited an increase in green fluorescence starting about 8 h after treatment initiation (Figure S4A), similar to the kinetic of the CalRexin™:pHrodo™ Red signal obtained with this reagent only (Figure 1A). Acquiring both green and red signal from these cells 24 hours after exposure to 8 μM rapamycin revealed two major populations: one exhibiting only green fluorescence (from CYTO-ID®) and one exhibiting both signals, suggestive of co-localization (Figure S4C). However, merged confocal images obtained at higher magnification largely point to be very limited co-localization of green and red puncta (Figure S4D & E), confirming that CYTO-ID® and CalRexin™:pHrodo™ Red stain different subcellular compartments.

Finally, sHeLa cells exposed to 8 μM rapamycin and stained with both CalRexin™:pHrodo™ Red and LysoTracker™ Green - an organic dye that accumulates in lysosomes driven by pH – appear to exhibit elevated fluorescence in both the green and red channels at low magnification (Fig. 2A-C). Higher magnification imaging revealed that in this conditions, most fluorescent organelles are either yellow (as stained by both CalRexin™:pHrodo™ Red and LysoTracker™ Green) or green only (as stained by LysoTracker™ Green), with only few red only organelles (as stained by CalRexin™:pHrodo™ Red) (Fig. 2D-L and Video). These data are in line with the ability of CalRexin™:pHrodo™ Red to stain autophagy-associated lysosomes but not other lysosomal compartments, which are instead not discriminated by LysoTracker™ Green.

**Figure 2.**
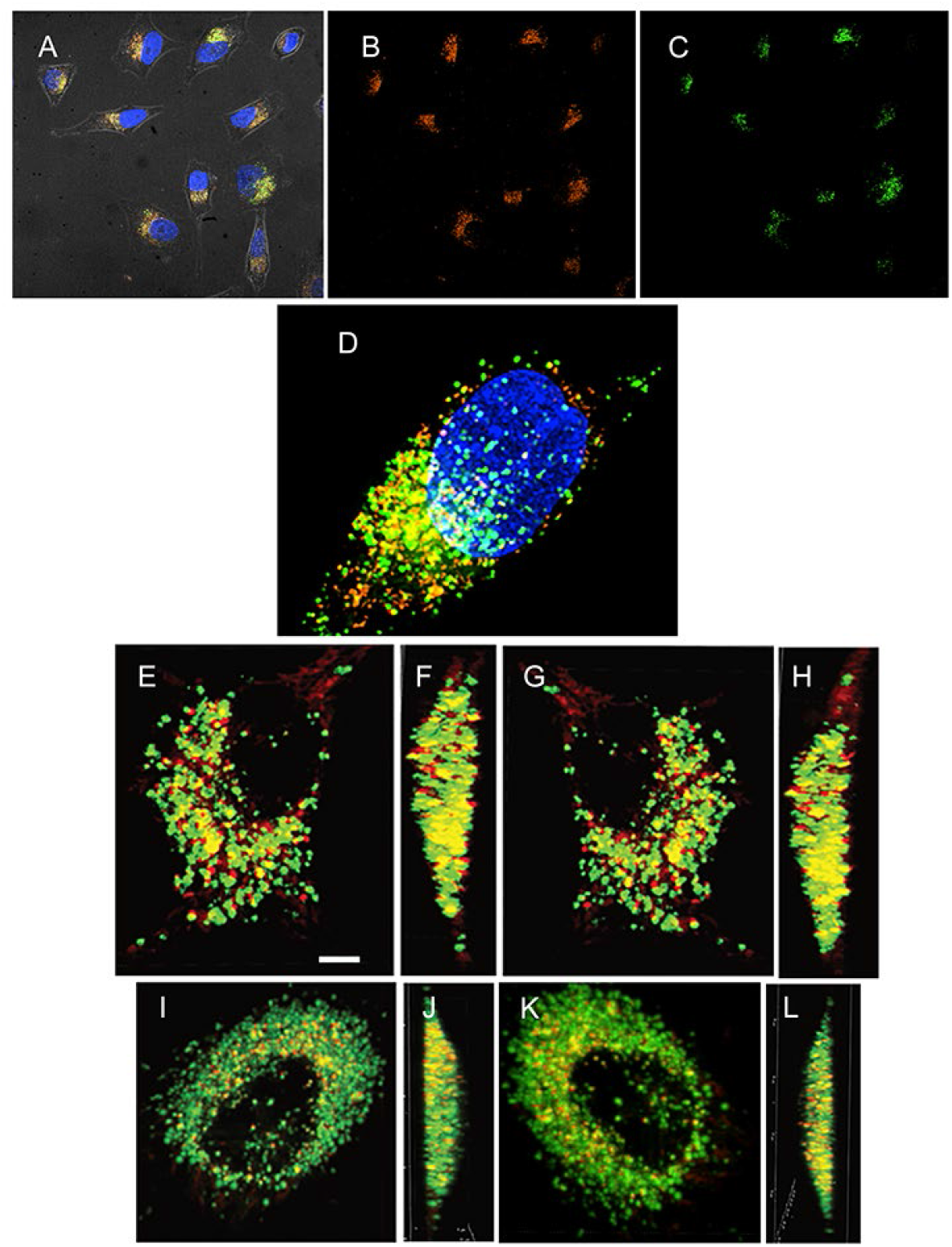
Images of live sHeLa cells treated with 8 μM rapamycin or FBS starvation for 24 hours. **Figure 2A-C**. Operetta images at 24 h: panel A, merged; panel B, CalRexin™:pHrodo Red; panel C, LysoTracker™ Green. **Figure 2D**. Representative, merged deconvoluted confocal image with CalRexin™:pHrodo Red and LysoTracker™ Green in a rapamycin treated sHeLa cell. **Figure 2E-H**. Single plane images of sHeLa cells treated with rapamycin for 24 h and imaged with CalRexin™:pHrodo™ Red and LysoTracker™ Green using the W1-SoRa super resolution spinning-disk mode. Panel E, overview of cell; F, sideview; G, underview of cell; H, alternative sideview of cell. **Figure 2I-L**. Single plane images of sHeLa cells starved of FBS for 24 hours and imaged with CalRexin™:pHrodo™ Red and LysoTracker™ Green. Panel I, overview of cell; J, sideview; K, underview of cell; L, alternative sideview of cell. The calibration bar in panel E is 50 μM in panels A-C and 5 μM in panels D-L.

## Discussion

Here, we provide experimental validation to the ability of the CT Receptor-targeted pH-sensitive fluorescent dye CalRexin™:pHrodo™ Red to enable high throughput/content imaging of autophagy as a potential approach to screen for novel autophagy modulators. This largely reflects the ability of CalRexin™:pHrodo™ Red to selectively accumulate in autophagy-associated lysosomes via late endosomes (amphisomes) as shown in the schematic Figure 3 – hence differing from CYTO®-ID, which mostly stains autophagosomes, and LysoTracker™ Green, which stains lysosomes based on pH, irrespective of source - and emit a bright and stable red fluorescence at an acidic pH. In line with this notion, sHeLa cells exposed to conventional autophagy activators such as rapamycin and serum starvation emitted a bright red signal that could be monitored over time by high throughput/content imaging, a signal that was lost when (1) cells were incubated with canonical autophagy inhibitors, (2) key autophagy genes were deleted, or (3) CalRexin™:pHrodo™ Red was replaced by an untargeted fluorescent isotype control.

**Figure 3.**
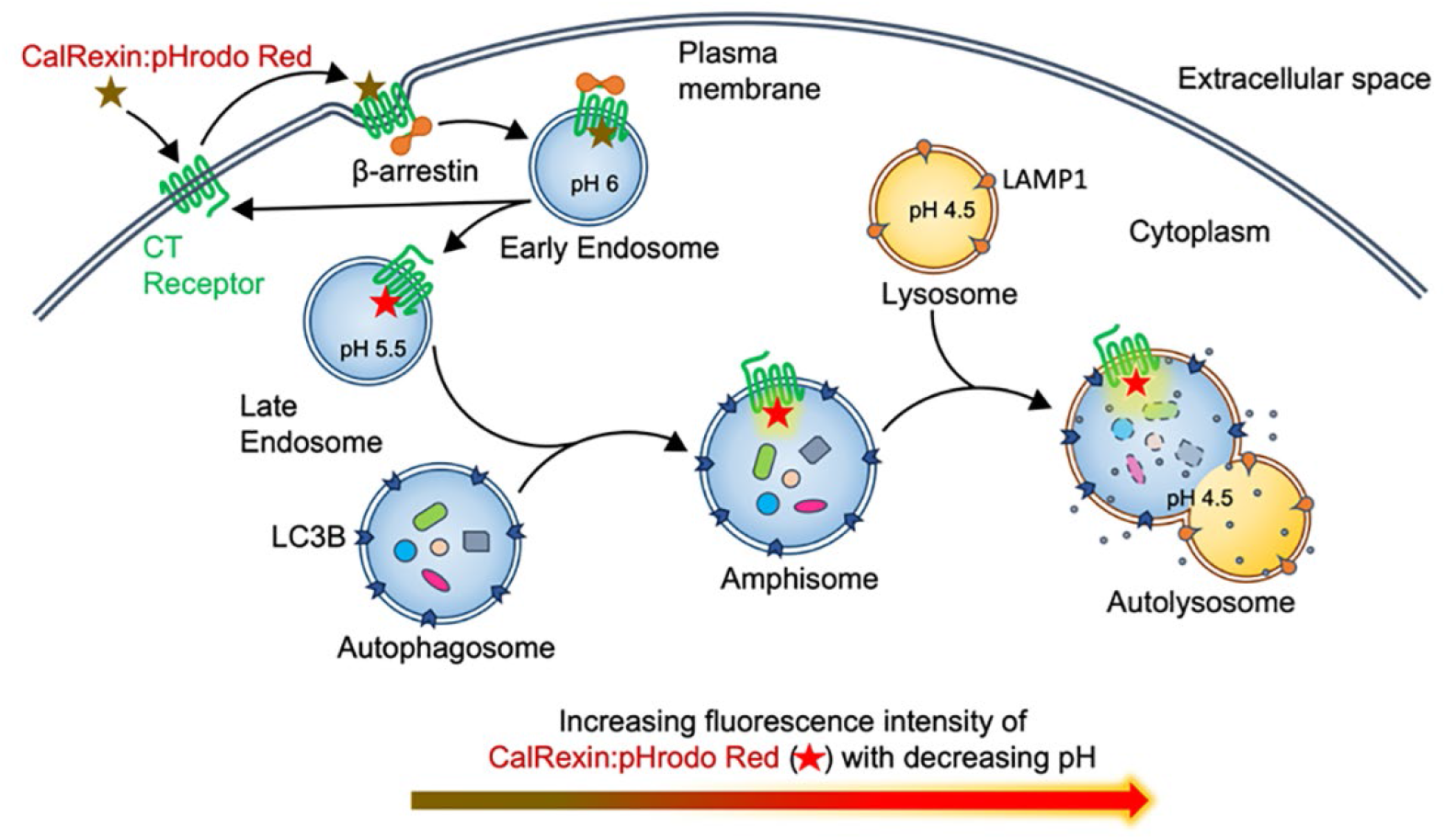
Schematic which represents the passage of CalRexin™:pHrodo™ Red bound to CT Receptor on the plasma membrane and is internalized through the early endosomes, late endosomes (pH 5.5) which fuse with the autophagosomes to form the amphisomes. The latter fuse with the lysosomes to form the autolysosomes (pH 4-5) which are identified with LysoTracker™ Green.

Cultured cells other than sHeLa exhibited similar signals, suggesting that CalRexin™:pHrodo™ Red is suitable to monitor autophagy across different cell types. We also tested this reagent with rapamycin induced autophagy in NIH 3T3 fibroblasts which resulted in reduced accumulation (data not shown) and most likely reflects lower affinity of the antibody for mouse CT Receptor. As a potential limitation, CalRexin™:pHrodo™ Red may not be suitable to monitor autophagy in cells lacking CT Receptor expression on the cell surface owing to genetic or epigenetic defects. How frequently healthy and malignant cells lose CT Receptor expression, however, remains to be determined. Along similar lines, the performance of CalRexin™:pHrodo™ Red may suffer from genetic or epigenetic defects compromising the coupling between autophagy and the endosome-amphisome pathway, but the frequency of such alterations is not clear. Despite these limitations, CalRexin™:pHrodo™ Red stands out as a promising dye for the implementation of high throughput/content screening efforts to identify novel modulators of autophagy.

## Supporting information

Ciccotosto et al Supplementary Material

## Acknowledgements

PG is supported by a postgraduate scholarship from the University of Melbourne. SGBF is funded as an Australian Research Council Future Fellow (FT180100543).

The LG lab is/has been supported (as a PI unless otherwise indicated) by one NIH R01 grant (#CA271915), by two Breakthrough Level 2 grants from the US DoD BCRP (#BC180476P1, #BC210945), by a grant from the STARR Cancer Consortium (#I16-0064), by a Transformative Breast Cancer Consortium Grant from the US DoD BCRP (#W81XWH2120034, PI: Formenti), by a U54 grant from NIH/NCI (#CA274291, PI: Deasy, Formenti, Weichselbaum), by the 2019 Laura Ziskin Prize in Translational Research (#ZP-6177, PI: Formenti) from the Stand Up to Cancer (SU2C), by a Mantle Cell Lymphoma Research Initiative (MCL-RI, PI: Chen-Kiang) grant from the Leukemia and Lymphoma Society [2], by a Rapid Response Grant from the Functional Genomics Initiative (New York, US), by a pre-SPORE grant (PI: Demaria, Formenti), a Collaborative Research Initiative Grant and a Clinical Trials Innovation Grant from the Sandra and Edward Meyer Cancer Center (New York, US), by startup funds from the Dept. of Radiation Oncology at Weill Cornell Medicine (New York, US), by industrial collaborations with Lytix Biopharma (Oslo, Norway), Promontory (New York, US) and Onxeo (Paris, France), as well as by donations from Promontory (New York, US), the Luke Heller TECPR2 Foundation (Boston, US), Sotio a.s. (Prague, Czech Republic), Lytix Biopharma (Oslo, Norway), Onxeo (Paris, France), Ricerchiamo (Brescia, Italy), and Noxopharm (Chatswood, Australia).

Our sincere gratitude is expressed for the role of Dr Tina Lavranos (former CEO) and Elpis Barons (current CEO) of Apop Biosciences for supporting this research project. We thank Dr Judy Scoble (CSIRO, Clayton) for organizing synthesis and purification of CalRexin™:pHrodo™ Red and Therapeutic Innovation Australia for supporting this work. We wish to acknowledge the contribution made by the Biological Optical Microscope Platform at the University of Melbourne for the provision and maintenance of the Operetta platform.

## Competing Interests

LG is/has been holding research contracts with Lytix Biopharma, Promontory and Onxeo, has received consulting/advisory honoraria from Boehringer Ingelheim, AstraZeneca, OmniSEQ, Onxeo, The Longevity Labs, Inzen, Imvax, Sotio, Promontory, Noxopharm, EduCom, and the Luke Heller TECPR2 Foundation, and holds Promontory stock options.

PJW is a director and owns stock in Apop Biosciences Pty Ltd that owns the anti-human CT Receptor antibody CalRexin™. PJW is also Chief Scientific Officer of Apop Biosciences Pty Ltd.

All other authors have no conflict of interest to declare.

## Author Contributions

GDC designed and executed the Operetta experiments, derived the data from images, organized the confocal microscopy and images were captured by JM. LG advised on the content in regard to autophagy and substantially re-drafted the manuscript. LM optimized the conjugation of CalRexin™:pHrodo™ Red for large batch synthesis. PG designed and drafted Figure 3 and commented on the draft manuscripts. SGBF suggested making CalRexin™:pHrodo™ Red in the first instance, contributed to the draft manuscript and co-supervised PG. ML provided the sHeLa Atg5 & FIP200 KO cell lines and contributed to the drafting of the manuscript. DLH provided support for the acquisition of data and supervised PG. PJW synthesised the first batch of CalRexin™:pHrodo™ Red, discovered the relationship between CalRexin™:pHrodo™ Red and autophagy, advised on the design of experiments, wrote multiple drafts of the manuscript and supervised PG. All authors have approved the final version and have agreed to be personally accountable for their own contributions.

## References

[1] Galluzzi L, Baehrecke EH, Ballabio A, et al. Molecular definitions of autophagy and related processes. EMBO J. 2017 Jul 3;36(13):1811–1836.

[2] Klionsky DJ, Abdel-Aziz AK, Abdelfatah S, et al. Guidelines for the use and interpretation of assays for monitoring autophagy (4th edition)(1). Autophagy. 2021 Jan;17(1):1–382.

[3] Sica V, Galluzzi L, Bravo-San Pedro JM, et al. Organelle-Specific Initiation of Autophagy. Mol Cell. 2015 Aug 20;59(4):522–39.

[4] de Morree A, Rando TA. Regulation of adult stem cell quiescence and its functions in the maintenance of tissue integrity. Nat Rev Mol Cell Biol. 2023 May;24(5):334–354.

[5] Debnath J, Gammoh N, Ryan KM. Autophagy and autophagy-related pathways in cancer. Nat Rev Mol Cell Biol. 2023 Aug;24(8):560–575.

[6] Kitada M, Koya D. Autophagy in metabolic disease and ageing. Nat Rev Endocrinol. 2021 Nov;17(11):647–661.

[7] Matsuzawa-Ishimoto Y, Hwang S, Cadwell K. Autophagy and Inflammation. Annu Rev Immunol. 2018 Apr 26;36:73–101.

[8] Sciarretta S, Maejima Y, Zablocki D, Sadoshima J. The Role of Autophagy in the Heart. Annu Rev Physiol. 2018 Feb 10;80:1–26.

[9] Klionsky DJ, Petroni G, Amaravadi RK, et al. Autophagy in major human diseases. EMBO J. 2021 Oct 1;40(19):e108863.

[10] Yamamoto H, Zhang S, Mizushima N. Autophagy genes in biology and disease. Nat Rev Genet. 2023 Jun;24(6):382–400.

[11] Galluzzi L, Green DR. Autophagy-Independent Functions of the Autophagy Machinery. Cell. 2019 Jun 13;177(7):1682–1699.

[12] Ganesan D, Cai Q. Understanding amphisomes. Biochem J. 2021 May 28;478(10):1959–1976.

[13] Kelly EE, Horgan CP, McCaffrey MW. Rab11 proteins in health and disease. Biochem Soc Trans. 2012 Dec 1;40(6):1360–7.

[14] Szatmari Z, Kis V, Lippai M, et al. Rab11 facilitates cross-talk between autophagy and endosomal pathway through regulation of Hook localization. Mol Biol Cell. 2014 Feb;25(4):522–31.

[15] Galluzzi L, Bravo-San Pedro JM, Levine B, et al. Pharmacological modulation of autophagy: therapeutic potential and persisting obstacles. Nat Rev Drug Discov. 2017 Jul;16(7):487–511.

[16] Levy JMM, Towers CG, Thorburn A. Targeting autophagy in cancer. Nat Rev Cancer. 2017 Sep;17(9):528–542.

[17] Stankov M, Panayotova-Dimitrova D, Leverkus M, et al. Flow cytometric analysis of autophagic activity with Cyto-ID staining in primary cells. Bio-protocol. 2014;4(7).

[18] Iwashita H, Sakurai HT, Nagahora N, et al. Small fluorescent molecules for monitoring autophagic flux. FEBS Lett. 2018 Feb;592(4):559–567.

[19] Hsu VW, Prekeris R. Transport at the recycling endosome. Curr Opin Cell Biol. 2010 Aug;22(4):528–34.

[20] Marchese A, Paing MM, Temple BR, Trejo J. G protein-coupled receptor sorting to endosomes and lysosomes. Annu Rev Pharmacol Toxicol. 2008;48:601–29.

[21] Huotari J, Helenius A. Endosome maturation. EMBO J. 2011 Aug 31;30(17):3481–500.

[22] Hu YB, Dammer EB, Ren RJ, Wang G. The endosomal-lysosomal system: from acidification and cargo sorting to neurodegeneration. Transl Neurodegener. 2015;4:18.

[23] Meeuwsen S, Persoon-Deen C, Bsibsi M, et al. tCytokine, chemokine and growth factor gene profiling of cultured human astrocytes after exposure to proinflammatory stimuli. Glia. 2003 Sep;43(3):243–53.

[24] Furness SGB, Hare DL, Kourakis A, et al. A novel ligand of calcitonin receptor reveals a potential new sensor that modulates programmed cell death [Article]. Cell Death Discovery. 2016 10/10/online;2:16062.

[25] Fukada S, Uezumi A, Ikemoto M, et al. Molecular signature of quiescent satellite cells in adult skeletal muscle. Stem Cells. 2007 Oct;25(10):2448–59.

[26] Gnocchi VF, White RB, Ono Y, et al. Further characterisation of the molecular signature of quiescent and activated mouse muscle satellite cells. PLoS One. 2009;4(4):e5205.

[27] Yamaguchi M, Watanabe Y, Ohtani T, et al. Calcitonin Receptor Signaling Inhibits Muscle Stem Cells from Escaping the Quiescent State and the Niche. Cell reports. 2015 Oct 13;13(2):302–14.

[28] Baghdadi MB, Castel D, Machado L, et al. Reciprocal signalling by Notch-Collagen V-CALCR retains muscle stem cells in their niche. Nature. 2018 May;557(7707):714–718.

[29] Marino G, Niso-Santano M, Baehrecke EH, Kroemer G. Self-consumption: the interplay of autophagy and apoptosis. Nat Rev Mol Cell Biol. 2014 Feb;15(2):81–94.

[30] Ma Y, Galluzzi L, Zitvogel L, Kroemer G. Autophagy and cellular immune responses. Immunity. 2013 Aug 22;39(2):211–27.

[31] Nguyen TN, Padman BS, Usher J, et al. Atg8 family LC3/GABARAP proteins are crucial for autophagosome-lysosome fusion but not autophagosome formation during PINK1/Parkin mitophagy and starvation. J Cell Biol. 2016 Dec 19;215(6):857–874.

[32] Nguyen TN, Lazarou M. A unifying model for the role of the ATG8 system in autophagy. J Cell Sci. 2022 Jun 1;135(11).

[33] Lazarou M, Sliter DA, Kane LA, et al. The ubiquitin kinase PINK1 recruits autophagy receptors to induce mitophagy. Nature. 2015 Aug 20;524(7565):309–314.

[34] Chen M, Ren X, Nguyen TN, et al. Structure and activation of the human autophagy-initiating ULK1C:PI3KC3-C1 supercomplex. bioRxiv. 2023:2023.06.01.543278.

[35] Wookey PJ, Gupta P, Bittencourt L, et al. Methods to measure calcitonin receptor activity, up-regulated in cell stress, apoptosis and autophagy. F1000Research. 2021;10.

[36] Kimura S, Noda T, Yoshimori T. Dissection of the autophagosome maturation process by a novel reporter protein, tandem fluorescent-tagged LC3. Autophagy. 2007 Sep-Oct;3(5):452–60.

[37] Hirano M, Ando R, Shimozono S, et al. A highly photostable and bright green fluorescent protein. Nat Biotechnol. 2022 Jul;40(7):1132–1142.

[38] Oeste CL, Seco E, Patton WF, et al. Interactions between autophagic and endo-lysosomal markers in endothelial cells. Histochem Cell Biol. 2013 May;139(5):659–70.

