## Supplementary material for "A novel pH-sensitive probe to quantify autophagy on high throughput/content imaging platforms": Ciccotosto et al Supplementary Material

**Methods**

**Cell lines**

The MG63 fibroblast cell line (ATCC #CRL-1427) derived from a human osteosarcoma and in this study was cultured in MEMα (TFS 12561056) and supplemented with 10% FBS. The mouse embryonic fibroblast NIH 3T3 cell line (ATCC #CRL-1658) was cultured in DMEM/F12 (TFS 11320-033) and supplemented with 10% (v/v) FBS, and 10 U/mL penicillin-streptomycin (TFS 15140122). Both cell lines were cultured in a humidified incubator with 5% $CO_2$ and 37$^oC$. Cell lines sHeLa WT, sHeLa *Atg5* and sHeLa *RB1CC1* were cultured as described in the Methods section.

Cells were maintained in uncoated T75 flasks in a humidified 37$^oC$ incubator supplemented with 5% $CO_2$. Cells were dislodged with Trypsin/EDTA, harvested, pelleted and a single cell suspension prepared and the number of cells counted. Cells were seeded at 10,000 and 20,000 cells per well in 96 well plates (black wall optically-clear cyclic olefin bottom plate (CellCarrier-96 Ultra, PerkinElmer #6055302)) and μ-Slide 8 well (DKSH #80821) slides respectively that were treated with Matrigel (Corning #354230*).* Experiments were performed using cell lines passaged less than 20 times. Cells were allowed to adhere to surface overnight before treatment and imaging.

**Figure S1.**

**Figure S1A.** In buffers prepared with a range of pH values, the fluorescence signal (normalized relative fluorescent units [RFU]) of CalRexin™:pHrodo™ Red is shown. CalRexin™:pHrodo™ Red fluoresces brightly at low pH.

**Figure S1B.** In the cell line MG63, increasing fluorescence of CalRexin™:pHrodo™ Red was measured over 24 hours on the Operetta platform and the response to a range of rapamycin concentrations (vehicle, open squares; 2 µM, closed inverted triangles; 4 µM, closed triangles; 8 µM, closed squares; 10 µM, closed diamonds).

**Figure S1C.** CalRexin™:pHrodo Red (4 µg/mL) [8 µM rapamycin, blue closed squares; vehicle, blue open squares] versus isotype control mouse IgG1:pHrodo Red (4 µg/mL) [8 µM rapamycin, closed orange squares; vehicle, open orange squares]. CR = CalRexin™:pHrodo™ Red; IC = IgG1 isotype control:pHrodo™ Red

**Figure S1D.** HeLA wildtype cells induced with 8 µM rapamycin versus vehicle controls. Comparison of the responses over 24 hours imaged with CalRexin™:pHrodo™ Red (batch 1, DoL 1.9, succinimidyl ester-pHrodo™ Red, TFS #P36600) [rapamycin, closed blue squares; vehicle, open blue squares] and CalRexin™:pHrodo™ Red (batch 2, DoL 1.9, iFL STP ester, TFS #P36010) [rapamycin, closed red squares; vehicle, open red squares].

Mean values of three independent experiments +/- SEM.

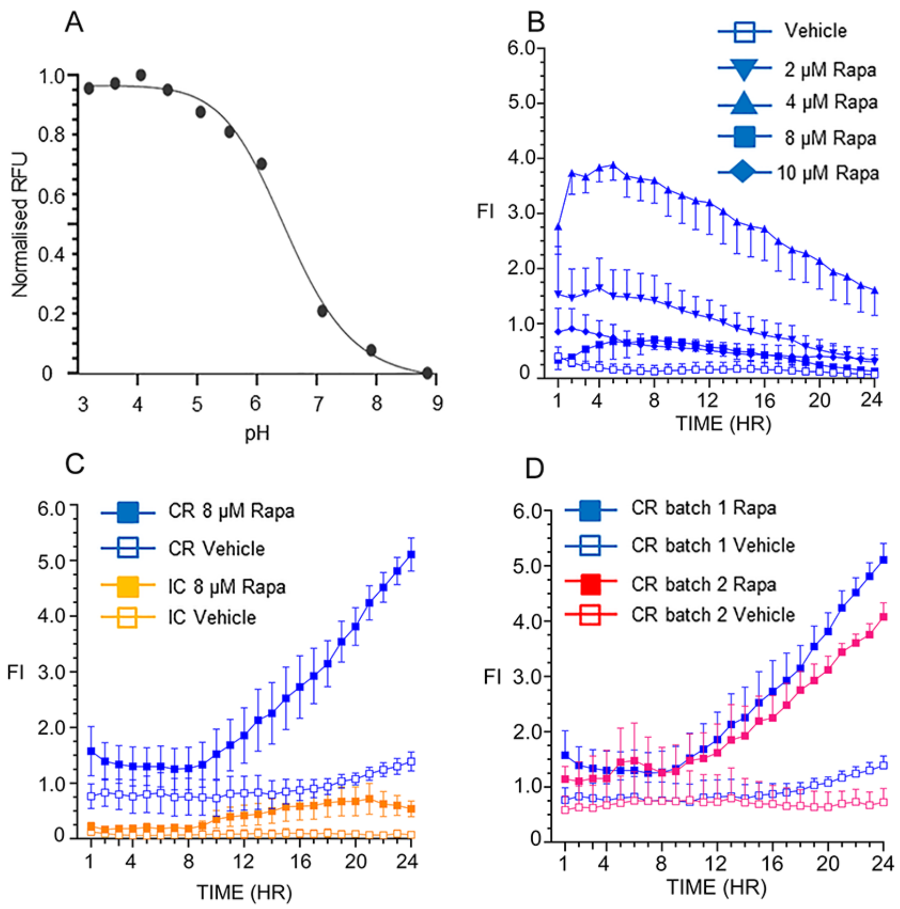

**Figure S2.** The effect of serum starvation (no FBS), depleted serum (5% dFBS & 5% eFBS) and serum replete (5% FBS) on the fluorescence signal of CalRexin™:pHrodo™ Red measured on the Operetta platform in several cell lines.

**Figure S2A.** (sHeLa): no FBS, blue closed squares; 5% eFBS, blue closed circles; 5% dFBS, blue closed diamonds; 5% FBS, blue open squares.

**Figure S2B.** (5% dFBS): sHela WT, closed blue diamonds; KO *Atg5*, closed red diamonds; KO *RB1CC1 (FIP200)*, closed green diamonds.

**Figure S2C.** (5% eFBS): sHela WT, closed blue circles; KO *Atg5*, closed red circles; KO *RB1CC1*, closed green circles.

Mean values of three independent experiments +/- SEM.

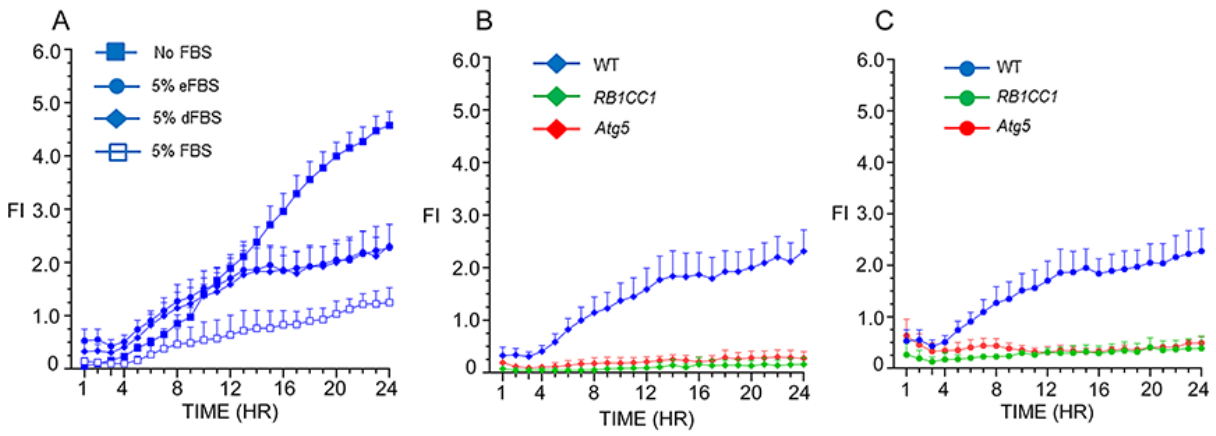

**Figure 3.** Transient transfected sHela cells (WT) with the Premo$^{TM}$ tandem sensor (RFP-GFP-LC3B) treated with 8 µM rapamycin and imaged on the Operetta over 24 h.
**Figure S3A.** The change in FI over 24 h for GFP: 8 µM rapamycin, blue closed squares; vehicle, blue open squares; 8 µM rapamycin and 50 µM chloroquine, blue closed inverted triangles; 50 µM chloroquine, blue open inverted triangles.
**Figure S3B.** The change in FI for RFP: 8 µM rapamycin, blue closed squares; vehicle, blue open squares; 8 µM rapamycin and 50 µM chloroquine, blue closed inverted triangles; 50 µM chloroquine, blue open inverted triangles.
**Figure S3C.** Shows the high number of cells expressing Premo$^{TM}$ tandem sensor (red channel) at 1 h.
**Figure S3D.** Shows RFP alone at 24 h.
**Figure S3E.** Shows co-localisation (yellow) of the tandem sensors, GFP-LC3B and RFP-LC3B at 24 hr.

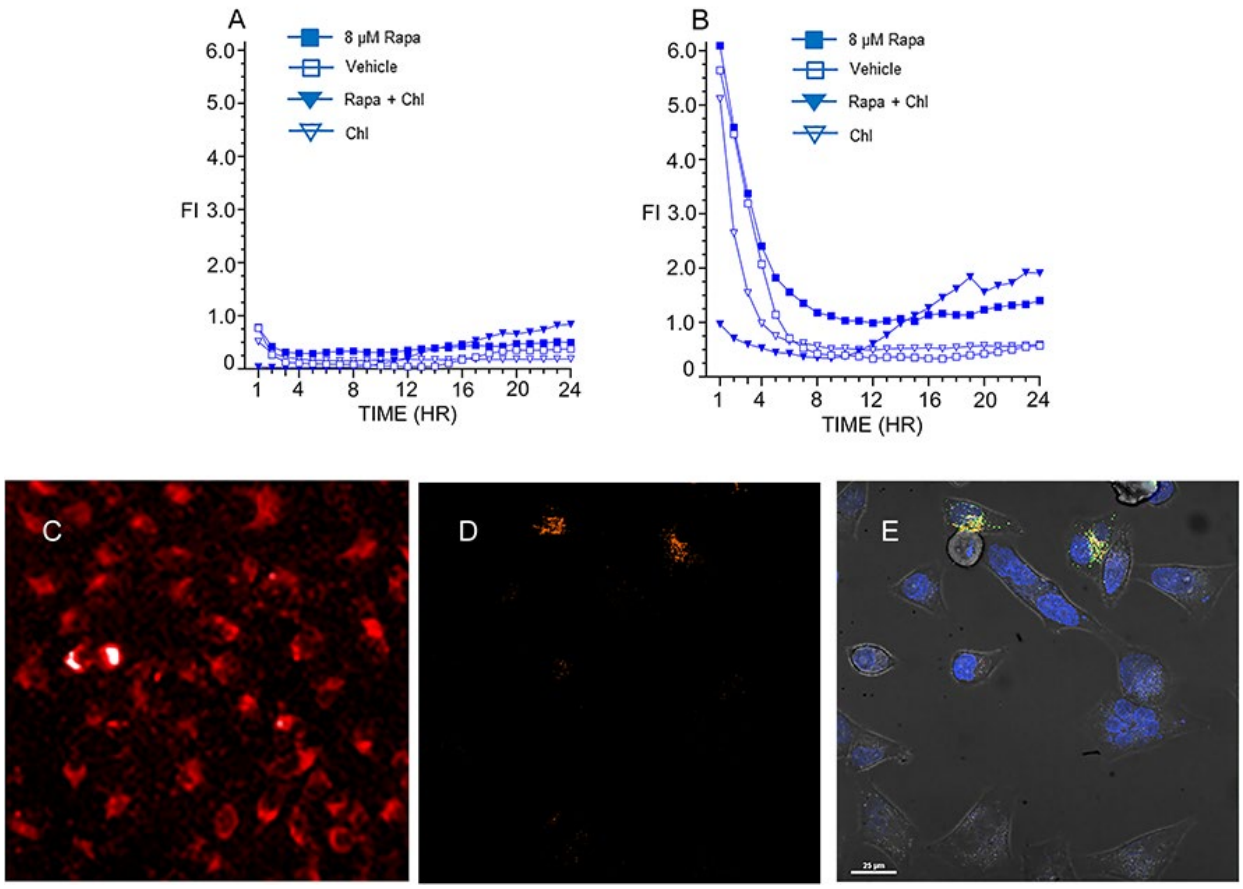

**Figure S4.** sHeLa cells untreated and treated with 8 µM rapamycin, and imaged with CYTO-ID® (ENZO #ENZ-kit175, green) and CalRexin™:pHrodo™ Red (Apop Biosciences, red).

**Figure S4A**. Accumulation of CYTO-ID® fluorescence intensity with rapamycin at 24 hours. The fluorescence intensity in this graph is measured in the green channel: 8 µM rapamycin, blue closed squares; vehicle, blue open squares; rapamycin plus chloroquine, blue closed inverted triangles; chloroquine alone, blue open inverted triangles. Mean values of three independent experiments +/- SEM.

**Figure S4B.** Operetta image of CYTO-ID® channel alone with 8 µM rapamycin at 24 hours.

**Figure S4C.** Operetta image of CYTO-ID® and CalRexin™:pHrodo™ Red staining with 8 µM rapamycin at 24 hours in which potential co-localisation is indicated in yellow.

**Figure S4D** & **E.** High magnification images of CYTO-ID® and CalRexin™:pHrodo™ Red staining with 8 µM rapamycin at 24 hours in which co-localisation is indicated in yellow.

The calibration bar in C is 25um in B & C and the bars in panels D & E are 10 µm.

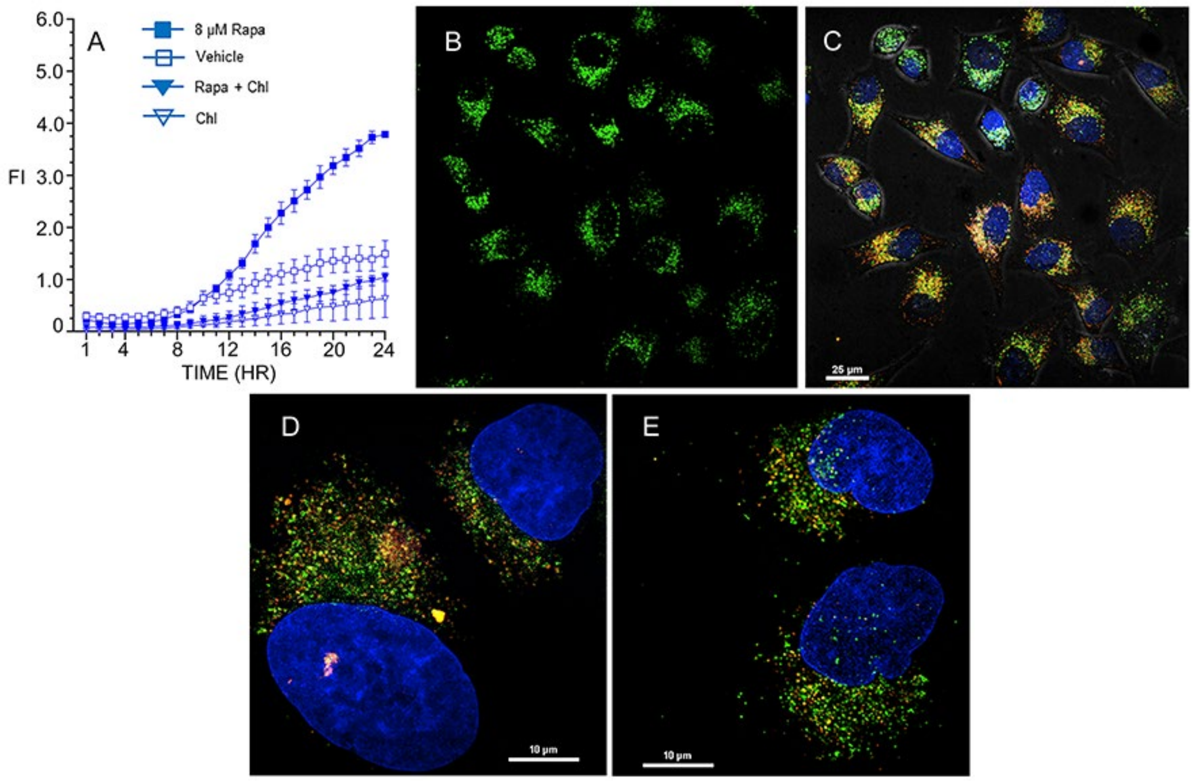

| |
|---|
| **Movie S5**. made of single sHeLa cells from Z-stacks of confocal images of live cells treated with 8 µM rapamycin for 24 hours and imaged on the Nikon CSA-W1 SoRa confocal microscope. CalRexin™:pHrodo™ Red and LysoTracker™ Green. |
| Movie<br>Rapaymycin treated for 24h wt sHela cells combined.avi |
